# Disentangling the effects of metabolic cost and accuracy on movement vigor

**DOI:** 10.1101/2023.02.08.527734

**Authors:** Garrick W. Bruening, Shruthi Sukumar, Robert J. Courter, Megan K. O’Brien, Alaa A. Ahmed

## Abstract

On any given day, we make countless reaching movements to objects around us. While such ubiquity may suggest uniformity, each movement is unique in the speed with which it is made. Some movements are slow, while others are fast. These variations in reach speed have long been known to be influenced by accuracy constraints; we slow down when accuracy demands are high. However, in other forms of movement like walking, metabolic cost is the primary determinant of movement speed. Here we ask, how do metabolic cost and accuracy interact to determine speed of reaching movements? First we systematically measure the effect of increasing mass on the metabolic cost of reaching across a range of movement speeds. Next, in a sequence of three experiments, we examine how added mass affects preferred movement speeds in a simple reaching task with increasing accuracy requirements. We find that mass consistently increased metabolic cost and led to slower movements. Yet, intriguingly, preferred reach speeds were slower than metabolically optimal. We then demonstrate how a single model that, critically, considers both accuracy and metabolic cost can explain preferred movement speeds across the range of accuracy and effort requirements tested. Together, these findings provide a unifying framework to explain the combined effects of metabolic cost and accuracy on movement speed, and also highlight the integral role metabolic cost plays in determining reach speed.

## Introduction

A defining characteristic of any movement is the speed, or *vigor*, with which it is performed. While more valuable reach targets elicit higher vigor Summerside et al. [2018], decreased reach vigor is often attributed to increased accuracy constraints associated with the target, mainly owing to the speed-accuracy tradeoff Fitts [1954]. Faster and quicker movements tend to be less accurate, leading to slower speed selection in the face of high accuracy demands. On the other hand, movements incur objective costs in the form of metabolic energy expenditure. For instance, in locomotion, metabolic energy minimization explains the selection of speed, step frequency and stride length Batliner et al. [2017], Ralston [1958], Selinger et al. [2015]. These kinematics are selected in a manner that minimizes effort when represented as metabolic cost per unit distance. The metabolic costs of reaching movements, however, remain unmeasured; instead, the effort cost of reaching has been estimated by biomechanical proxies such as derivative of acceleration, or sum of squared torque. Yet recent evidence suggests reaching movements are likewise sensitive to metabolic costs Shadmehr et al. [2016]. For example, we reach slower in directions that involve moving more mass and prefer to reach in directions of lower effort Goble et al. [2007], Gordon et al. [1994]. While these data directly implicate metabolic cost in the selection of reach vigor, this modulation of reaching effort has not been confirmed through collection of metabolic expenditure data. Furthermore, it is unclear how reach vigor is modulated in the presence of increased demands on both accuracy and effort.

The primary aim of this work was to provide a comprehensive examination into the interacting roles of effort and accuracy in control of reach vigor. To this end, we first measured metabolic costs of reaching movements, as a representation of effort, and demonstrated that metabolic costs increase with both speed and the mass required to transport at the hand. Secondly, we quantified preferred reaching speeds in an array of loading conditions and accuracy constraints and established that self-selected speeds become slower when effort costs increase, even when controlling for accuracy Fitts [1992]. Next we used a neuroeconomic modeling approach to determine the combined influence of metabolic and accuracy costs on the selection of movement speed, or vigor. We found that maximization of net reward rate, which considered metabolic cost and accuracy, provided excellent predictions of preferred movement speed. Interestingly, minimizing metabolic cost alone could not fully capture preferred movement speeds. These results highlight that metabolic energy unequivocally influences the selection of arm reaching vigor, and together with accuracy and time, provide evidence for a neuroeconomic framework based on net reward rate that determines movement speed.

## Results

Our first step towards understanding the effect of effort on preferred movement speed was to quantify it. To do so, we considered metabolic cost as an objective representation of effort and measured these costs of reaching while changing speed and mass at the hand.

### The effect of mass on the metabolic cost of reaching

Healthy, young participants (N = 8) made 10 cm reaching movements at six prescribed speeds (ranging from Very, Very Slow [VS, 1.25-1.35 s] to Very, Very Fast [VVF, 0.225-0.275 s]) with four different masses (0 kg, 2.3 kg, 4.5 kg, and 9.1 kg) added at the hand for a total of 24 sets of reaching conditions (Fig. 1A). Conditions were blocked such that each consisted of five minutes of reaching (∼200 trials). As they performed the task, we measured metabolic rate via expired gas analysis. Metabolic rate was calculated at steady state from the final three minutes of reaching within each block.

**Figure 1:**
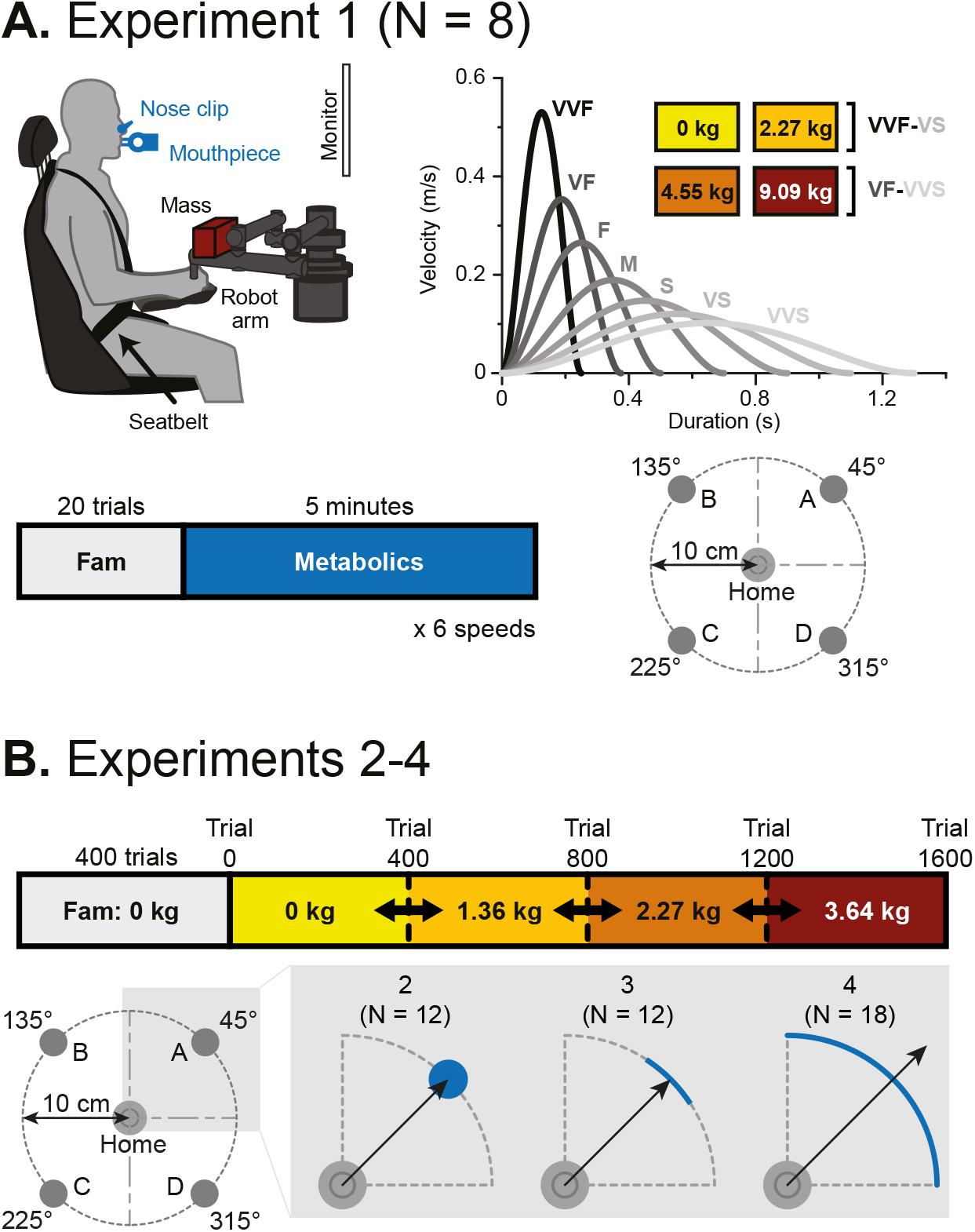
Experimental setup. (A) Experiment 1. *Top left*. Subjects made horizontal planar reaching movements while breathing into a mouthpiece. Mass was added at the hand. *Bottom right*. Subjects made out-then-back reaching movements across a range of added masses and speeds to four targets 10 cm from the home circle. *Top right*. There were seven distinct speeds, and subjects completed six of those speeds within each mass. The two heaviest masses corresponded with the six slowest speeds: Very Fast (VF, 0.325-0.425 s) to Very, Very Slow (VVS, 1.25-1.35 s). The two lighter masses corresponded with the six faster speeds: Very, Very Fast (VVF, 0.225-0.275 s) to Very Slow (VS, 1.05–1.15). *Bottom left*. The number of trials within each speed was set to allow for approximately five minutes of reaching. (B) Experiments 2-4. *Top row*. Subjects underwent five blocks of reaching movements including a familiarization block and four added mass blocks. The order of mass conditions was randomized for each subject. The general setup is the same as experiment 1. Subjects completed 400 trials in familiarization and each mass condition. *Bottom row*. Subjects made goal-directed reaching movements from a home circle and stopped at one of four targets, 10 cm away from the home circle. In experiment 2, subjects needed to stop in a circular target of the same size as experiment 1. In experiment 3, subjects needed to stop in a smaller, thin, arc-shaped target. In experiment 4, subjects did not need to stop and performed out-and-back reaching movements, with the only accuracy criteria that they cross the perimeter of the outer circle within the 90 degree quadrant.

#### Metabolic expenditure increases with added mass under varying speed conditions

Before participants performed the reaching task, we measured their resting metabolic rate, *ė*_*r*_ in three five-minute baseline periods, as they sat quietly in the experimental chair. On average, the resting metabolic power was *ė*_*r*_ = 73.33 ± 3.6 W. As they performed the reaching task, gross metabolic power increased with faster reaching speeds (*β* = -7.67e-1, p *<* 2e-16; Fig. 2A); with no added mass, gross metabolic power ranged from 92.78 ± 6.63 W for the slowest reach to 171.98 ± 18.51 W for the fastest reach. Furthermore, across movement speeds, gross metabolic power increased significantly with added mass (*β* = 1.73e-2, p = 2.52e-7; Fig. 2A). For a movement at the second fastest speed condition, adding 9.1kg of mass at the hand led to an increase in gross metabolic power from 131.65 ± 14.32 W to 222.06 ± 24.39 W, a nearly 70% increase. Thus, faster reaches and greater mass both led to increased metabolic expenditure.

**Figure 2:**
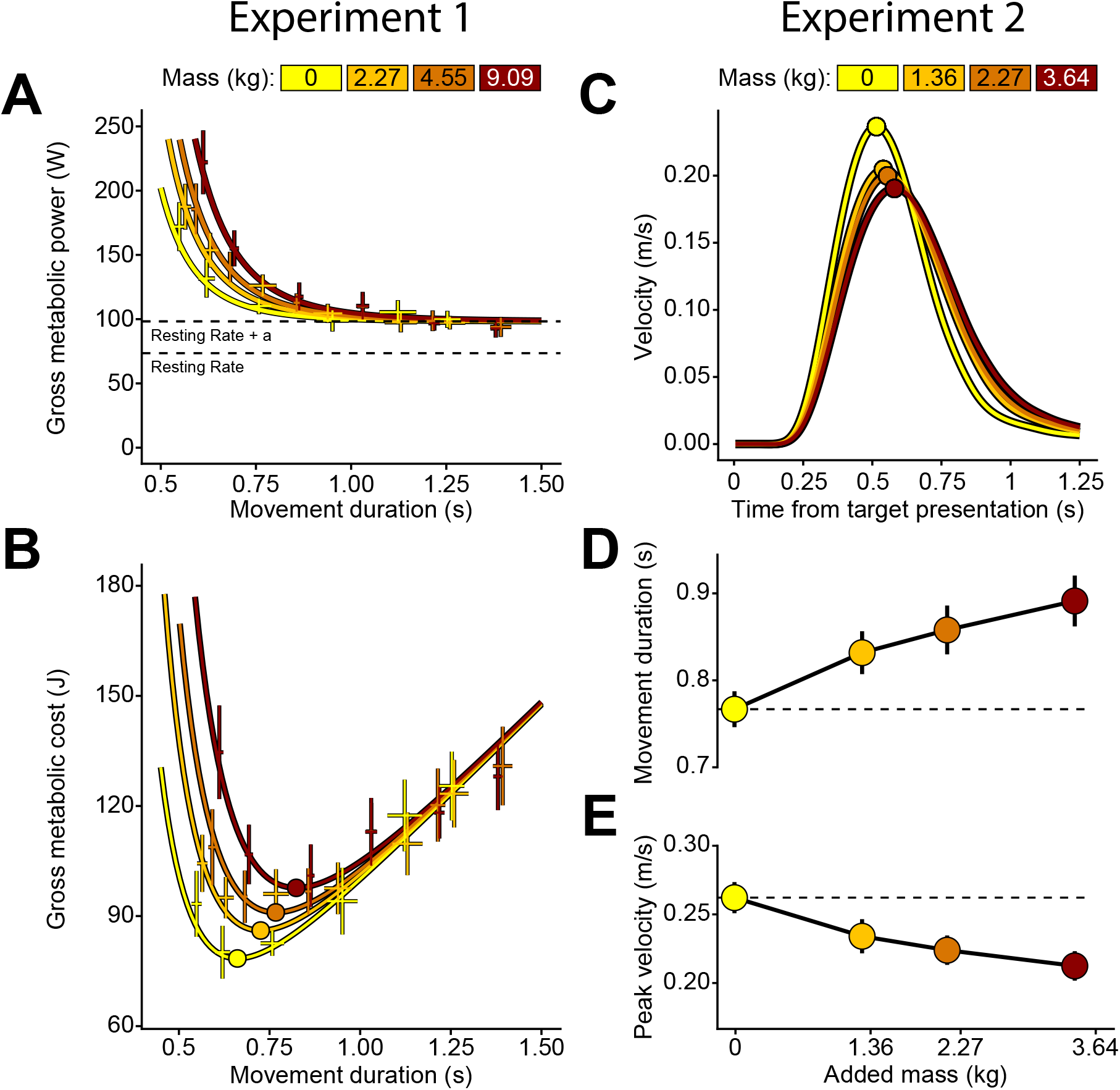
*Left column*. Experiment 1 results. (A) Gross metabolic power increases with added mass and movement speed (shorter durations). (B) Gross metabolic cost shows a distinct minimum, and this minimum duration increases with added mass. *Right column*. Experiment 2 results. (C) Average velocity traces per each added mass condition. Each line represents the subject average for one mass condition, with the peak velocity indicated by a circle. In panels D and E, each point represents the subject average for a specific mass condition. Horizontal dashed lines show the average value for 0 kg of added mass. (D) Mass increased movement duration. (E) Mass reduced peak velocity. Error bars are standard error of the mean across subjects.

We next sought to determine how mass influenced the metabolically optimal movement speed, i.e., the speed at which the metabolic cost of the movement was at a minimum. To do so, we parameterized metabolic rate as a function of mass and movement duration by fitting the gross metabolic rate data to the following equation based upon the observed effects of mass on the metabolic cost of walking (equation 1). In equation 1, *m* represents the effective mass of the arm with the added mass (see Methods), and *t*_*m*_ is the movement duration. The best fit parameters were *a* = 98.25 ± 3.05, *b* = 0.86 ± 0.43, *i* = 0.83 ± 0.10, and *j* = 5.83 ± 0.60 (SSE = 120872.75, AIC = 1750.83).

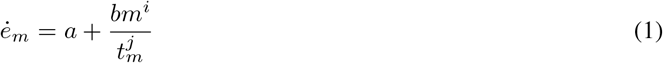

We also tested an alternative formulation wherein the constant, *a*, was also scaled by the effective mass of the movement raised to a power (equation 2). In equation 2, the fitted value for *k* was not statistically different from zero and the model performed similarly compared to equation 1 (*k* = -0.02 +/-0.04, SSE = 120743.07, AIC = 1752.63), indicating that the time-invariant component of metabolic power did not change with added mass. Thus, we moved forward with equation 1.

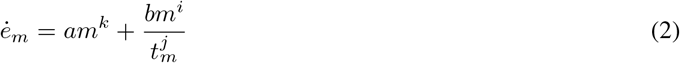

To obtain the total metabolic cost of that reach in joules, metabolic rate (*e*_*m*_, *ė*_*m*_) measured in (Joules/s) in equation 1 is multiplied by the movement duration *t*_*m*_, giving us metabolic cost in equation 3.

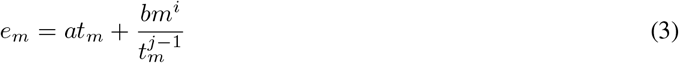

Indeed, the empirical data revealed that metabolic cost of the fastest reaches were high, reducing as the movement slows down, but then increasing again at slower speeds (Figure 2B). The minima of these curves represents the movement duration that minimizes the metabolic cost of the reach. The movement duration that minimized metabolic cost in equation 3 at increasing load conditions 0, 2.27, 4.55, and 9.09 kg were 0.66 s, 0.73 s, 0.77 s, and 0.83 s, respectively. The corresponding metabolic cost at these movement durations were 78.61 J, 86.32 J, 91.24 J, and 97.92 J, respectively. We saw that not only does the metabolic minimum shift to longer movement durations at higher added load conditions, but the metabolic minimum cost increases with added mass. Therefore, the data from experiment 1 revealed that it was more energetically costly to make reaching movements with greater mass and that the metabolically optimal movement duration increased with added mass. In other words, metabolic energy minimization predicted slower movements with added mass.

### Added mass led to longer self-selected movement durations

Given that added mass leads to an increase in the metabolic cost of reaching, and that the metabolically optimal duration increases with added mass, we asked how this increased effort cost would influence an individual’s preferred movement speed. A second experiment was therefore designed to answer this question. Similar to experiment 1, participants made goal-directed reaching movements with added mass at the hand (Fig. 1B). However, instead of reaching at a prescribed speed, participants were free to reach at a self-selected speed.

At the beginning of each trial, one of four targets would appear centered on the circumference of a 10cm circle and participants were asked to reach and stop in the target to complete the trial. The experiment consisted of 1600 total trials, divided into four sets of 400 trials each. In each set of 400 trials, a different mass was added to the robot handle. The amount of added mass was hidden from the participant using an opaque container and could be 0 kg, 1.36 kg (3lbs), 2.27 kg (5lbs) or 3.64 kg (8lbs) (Fig. 1B). All participants experienced each mass condition once, and the order was randomized across participants.

Added mass led participants to make significantly slower movements as evidenced by significantly longer movement durations and lower peak velocities (Movement Duration: *β* = 3.29e-2, p *<* 2e-16, Peak Velocity: *β* = -1.31e-2, p *<* 2e-16; Fig. 2C-E). All else equal, when added mass at the arm was increased, individuals consistently opted for a slower movement speed. Therefore, not only do changes in added mass result in modulation of the energetics associated with the movement, but these changes in metabolic cost appear to influence the selection of reach speed.

### Predicting preferred movement duration

In experiments 1 and 2, we observed that mass increases the effort cost of movement and that subjects prefer slower movement durations for greater added mass. Can we explain these effort-based changes in movement preference in the context of a movement utility that is conserved across individuals? As a first step, we looked to what metabolic minimization would predict. For the mass and movement distances used in experiment 2, we calculated the metabolically optimal durations using Eq. 3. We found that the minimum metabolic durations were 0.66, 0.70, 0.73, and 0.75 s for the 0, 1.36, 2.27, and 3.64 kg conditions, respectively. Critically, the durations that minimized metabolic cost were much faster than the actual durations seen in experiment 2 (0.78, 0.84, 0.87, and 0.90 s for the 0, 1.36, 2.27, and 3.64 kg conditions, respectively). Even the metabolically optimal durations predicted from experiment 1, in which the added masses were larger, were faster than the observed durations from experiment 2 (0.66, 0.73, 0.77, and 0.83 s for the 0, 2.27, 4.55, and 9.09 kg conditions, respectively). While metabolic cost minimization correctly predicted a general slowing of movement, it could not fully explain preferred movement durations. These data suggest that additional factors beyond effort are being accounted for when prospectively optimizing movement speed.

#### Maximization of net reward rate explains the combined effect of effort and accuracy costs on preferred movement duration

If we assume that movements are made to optimize a certain objective, how does the metabolic cost of a reaching movement modulate the objective function optimized by the movement? We know that other factors, like reward and accuracy constraints of a movement, also affect the selected speed Summerside et al. [2018], Fick [1882]. Assuming that the purpose of movement is to successfully acquire reward as quickly as possible but with minimal effort, we can define the utility of movement, *J*, as the expected reward minus the associated costs of the movements, all divided by the total time taken to acquire the reward:

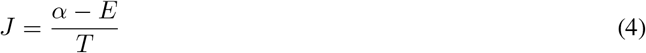

where *α* is the reward to be obtained, *E* is the effort cost associated with obtaining the reward, and *T* is the total time spent acquiring the reward. Substituting the terms in equation 4 specific to reaching movements, we represent the movement effort as measured metabolic cost of a reach with a given mass and duration, *e*_*m*_ (equation 3). Additionally, there is also the time spent preparing the movement, i.e., the reaction time. The effort cost associated with reaction time, *e*_*r*_, is equal to the metabolic expenditure when not reaching - the resting metabolic rate; this cost is represented as *e*?_*r*_ multiplied by the reaction time, *t*_*r*_, shown in equation 5. The total time to reward, *T*, then is the sum of the reaction time (*t*_*r*_) and movement duration (*t*_A*m*_). Finally, we have the expected reward, which is given by the probability of completing the movement successfully at a given *P* (*α*|*t*_*m*_, *m*), multiplied by the reward *α*. Therefore, we obtain our final utility function for completing a movement given by equation 5.

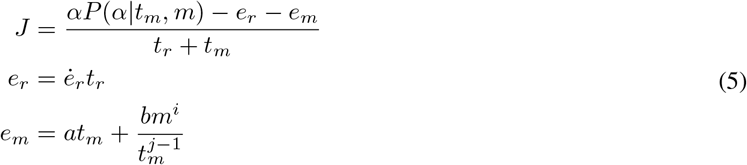

Critically, *P* (*α*|*t*_*m*_, *m*) represents the speed-accuracy tradeoff Fitts [1992]; the shorter the duration of the movement, the lower the probability of accurately completing it. This probability was modelled as a logistic function of movement duration and added mass (equation 6). To obtain empirical estimates of the parameters of this function, we fit the function to endpoint accuracy data from experiment 1 to capture the relationship between movement duration, mass, and the probability of successful movement completion. Successful movements in experiment 1 are those that end in the target of radius 1.4 cm. Any movement that ends outside is considered unsuccessful.

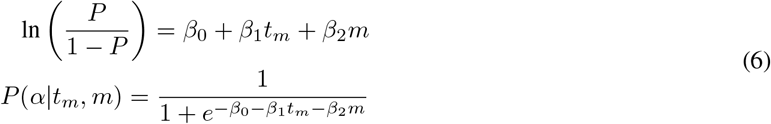

Figure 3A shows the fit to the probabilities measured from the data in experiment 1 to the corresponding parameters in equation 6 (*β*_0_= -1.45 ± 0.13, *β*_1_ = 5.88 ± 0.19, *β*_2_ = -0.10 ± 0.01). Using these parameters for *P* (*α*|*t*_*m*_, *m*) and the measured reaction times from experiment 2, we then used the net reward rate model in equation 5 to predict the selected movement durations in the data from experiment 2, fitting reward *α* as a free parameter. We found that net reward rate described preferred movement duration better than just metabolic cost alone (Figure 3B, *α* = 70.621). The predicted movement durations for experiment 2 were 0.758, 0.787, 0.808, and 0.835s for the added mass conditions 0, 1.36, 2.27, and 3.63 kg, respectively.

**Figure 3:**
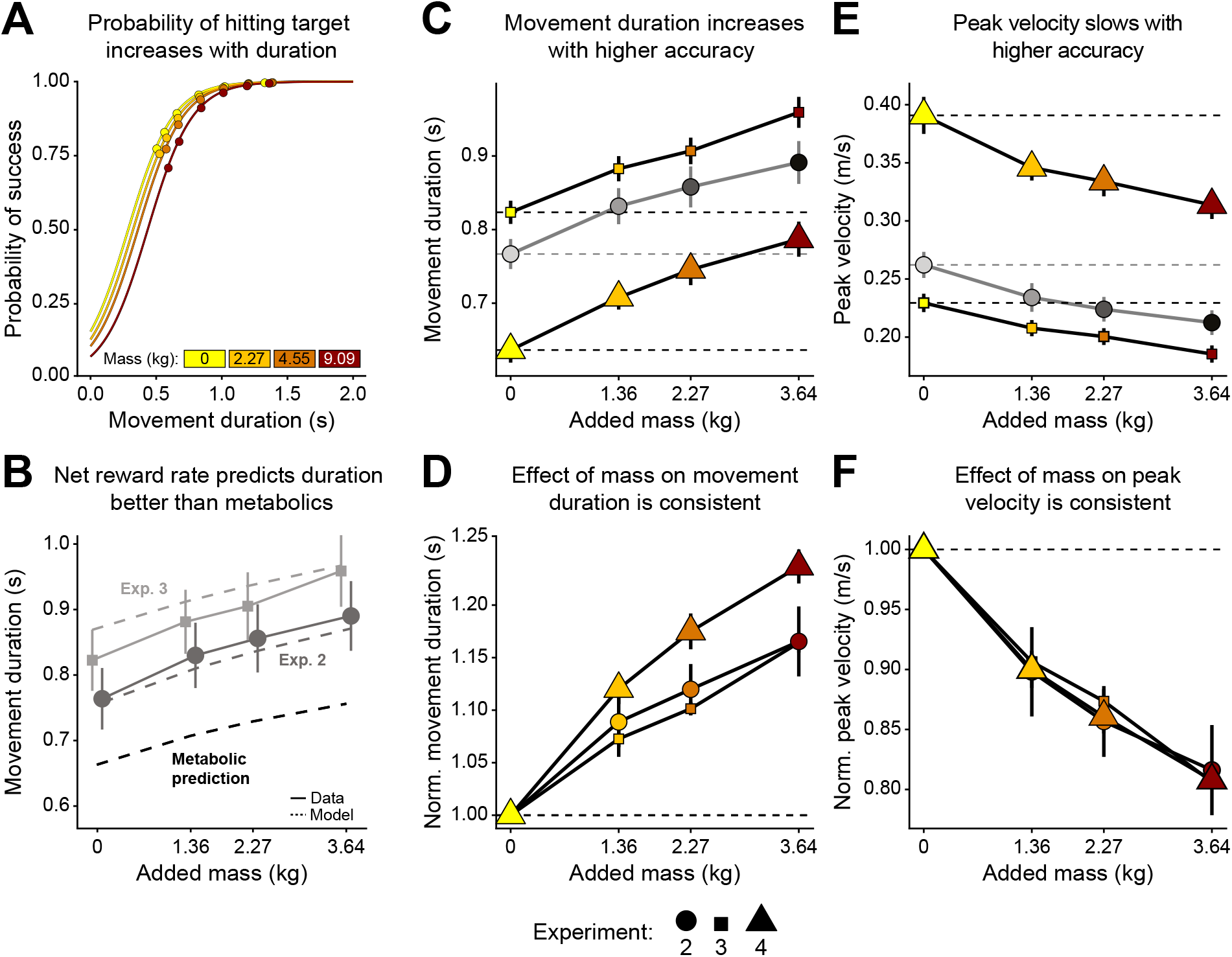
*Left column*. Modeling. (A) Function for the speed-accuracy trade off. Using the data from experiment 1, and the logistic regression shown in equation 6, we can compute the probability of success based on target accuracy. Data points are the fraction of trials within that condition that were a success. Each line is colored by mass added from experiment 1. (B) Optimal movement durations obtained by maximization of net reward rate across both experiments 2 and 3 using equation 5. Solid lines represent the average movement duration for experiment 2 (dark gray) and 3 (light gray). Error bars are standard error of subject means. Dashed lines indicate predicted optimal movement durations and are color-coded by experiment. The black dashed line indicates durations predicted by minimizing metabolic cost alone. *Middle and right columns*. Experiment 3 and 4 results. In all panels, each point represents the subject average for a specific mass condition. Error bars show standard error across subject averages. Horizontal dashed lines show the average value for 0 kg of added mass. Previously reported experiment 2 results are presented in gray (Fig. 2). (C) Mass and higher accuracy constraints increased movement duration. (E) Mass and higher accuracy constraints reduced peak velocity. Panels D and F: Movement metrics normalized as a fraction of each subjects 0 kg condition. (D) Movement duration and (E) peak velocity exhibited similar changes due to mass regardless of accuracy requirements.

In summary, we found that mass increased the metabolic cost of reaching and led to slower preferred movement speeds. These preferred movement durations could not be fully explained by metabolic cost minimization alone but were best explained as the outcome of decision aimed at maximizing the net reward rate of the movement, where net reward reflects the total reward to be acquired, the probability of acquiring that reward, and the metabolic cost of the movement.

### Controlling for accuracy of movement completion

Our results thus far demonstrate that increasing metabolic cost exerts a substantial influence on the selection movement speed. What remains unclear however is how effort and accuracy interacted to determine movement speed. Maximization of net reward rate assumes that effort and accuracy costs sum linearly to exert their combined influence on movement duration. However, it is possible that the effect of effort on duration may be mitigated (or augmented) when greater accuracy is demanded. To investigate this we ran an additional experiment, experiment 3, in which the accuracy requirements of the movement were increased by changing the success criterion for completing the movement. We reduced the target to a thin arc of 7 degrees with a thickness of 1 cm within which subjects were required to stop (Fig. 1B); this experiment represented a more stringent criterion than experiment 2. Thus we titrated the effect of accuracy on movement speed while keeping the modulation of added mass the same.

As expected, there was an effect of an experiment’s accuracy requirements on the preferred movement duration (ANOVA, p = 8.85e-5) and peak velocity (ANOVA, p *<* 1.17e-7) (Fig. 3C,E). Experiment 3, with its smaller target, exhibited longer movement durations and slower peak velocities. Yet even with the increased constraint on accuracy, movements slowed with added mass. Interestingly, the changes in movement duration due to mass were similar in extent to that observed in experiment 2 (exp 3: *β* = 3.53e-2, p *<* 2e-16; exp 2: *β* = 3.29e-2, p *<* 2e-16; Fig. 3D). Additionally, added mass influenced peak velocity similarly across both experiments (exp 3: *β* = -1.13 e-2, p *<* 2e-16; exp 2: *β* = -1.31e-2, p *<* 2e-16; Fig. 3F). Thus, the effect of effort on movement duration was conserved across accuracy constraints.

We next returned to the net reward rate model defined in equation 5 to investigate whether it could explain the combined effects of accuracy and effort on movement duration. The speed-accuracy tradeoff defined in equation 6 was first re-calculated using accuracy constraints that reflected those in experiment 3. Taking endpoint accuracy from experiment 1, we redefined a successful/accurate reach as one that instead finished within a 1 cm wide arc of 7 degrees (2.44 cm arc length). This resulted in a new set of parameters for the speed-accuracy tradeoff (*β*_0_ = -2.93 ± 0.09, *β*_1_ = 6.09 ± 0.13, *β*_2_ = -0.08 ± 0.01). Using these new accuracy parameters and the same reward *α* value fit from experiment 2, we once again used the net reward rate model to predict selected movement durations. Remarkably, we found that, without fitting any parameters to the observed movement durations in experiment 3, the net reward rate model nonetheless captured the data very well (Figure 3B).

While these findings indicated that accuracy and metabolic cost each made substantial, yet independent, contributions to movement speed, the possibility that the increased mass led to either real or perceived reductions in accuracy remained. To address this, we conducted a fourth and final experiment, where we relaxed the criterion for movement success. Subjects were only required to move the cursor through the quadrant and reach back to the home circle, and visual feedback of the cursor was removed. The purpose of this last experiment was to determine the effect of effort in the absence of any accuracy costs, to the extent that this possible. Importantly, we did not seek to compare movement durations to the previous experiments, as the movements themselves were different (out-and-stop vs. out-and-back). Our focus was on the effect of mass on within-experiment changes in movement speed. With accuracy costs removed, movement durations still slowed with increasing effort and the extent of change was similar to those observed in experiments 2 and 3 (exp 3: *β* = 3.53e-2, p *<* 2e-16; exp 4: *β* = 3.92e-2, p *<* 2e-16, exp 2: *β* = 3.29e-2, p *<* 2e-16; 3D). This is more clearly shown when comparing the normalized changes in movement duration with added mass, relative to the 0 mass condition across experiments (table 1). Added mass also slowed peak velocity of the reaching movements to the same extent across all three experiments (exp 3: *β* = -1.13 e-2, p *<* 2e-16; exp 4: *β* = -1.99e-2, p *<* 2e-16; exp 2: *β* = -1.31e-2, p *<* 2e-16; 3F). Even when accuracy costs were negligible, mass lead to a significant increase in movement duration and reduction in speed. This suggests that the changes in movement duration are driven by mass-related increases in effort, and not driven by mass-related changes in accuracy.

**Table 1:** Normalized movement durations for experiments 2, 3, 4, and metabolically optimal durations.

| Mass | Exp 2 | Exp 3 | Exp 4 | Metabolic optimum |
| --- | --- | --- | --- | --- |
| 0 kg | 1 | 1 | 1 | 1 |
| 1.36 kg | 1.086 | 1.071 | 1.114 | 1.065 |
| 2.27 kg | 1.121 | 1.100 | 1.174 | 1.098 |
| 3.64 kg | 1.164 | 1.166 | 1.237 | 1.140 |

In summation, these three experiments reveal that the qualitative relationship between metabolic cost and movement duration are preserved across accuracy conditions. This indicates that both the accuracy and effort of a movement significantly and independently modulate the selection of movement duration.

### Effort increased reaction time

An interesting finding across all experiments was that reaction time was chosen by the subject was also significantly modulated by the added mass condition. In experiment 1, while movement speed was restricted, reaction time was still freely chosen. Added mass led to an increase in self-selected reaction time (*β* = 2.40e-3, p *<* 2e-16; Fig. 4A). In experiments 2, 3, and 4 when movement speed was also self-selected, we found that this relation between reaction time and added mass was preserved (*β* = 4.47e-3, p *<* 2e-16; Fig. 4B).

**Figure 4:**
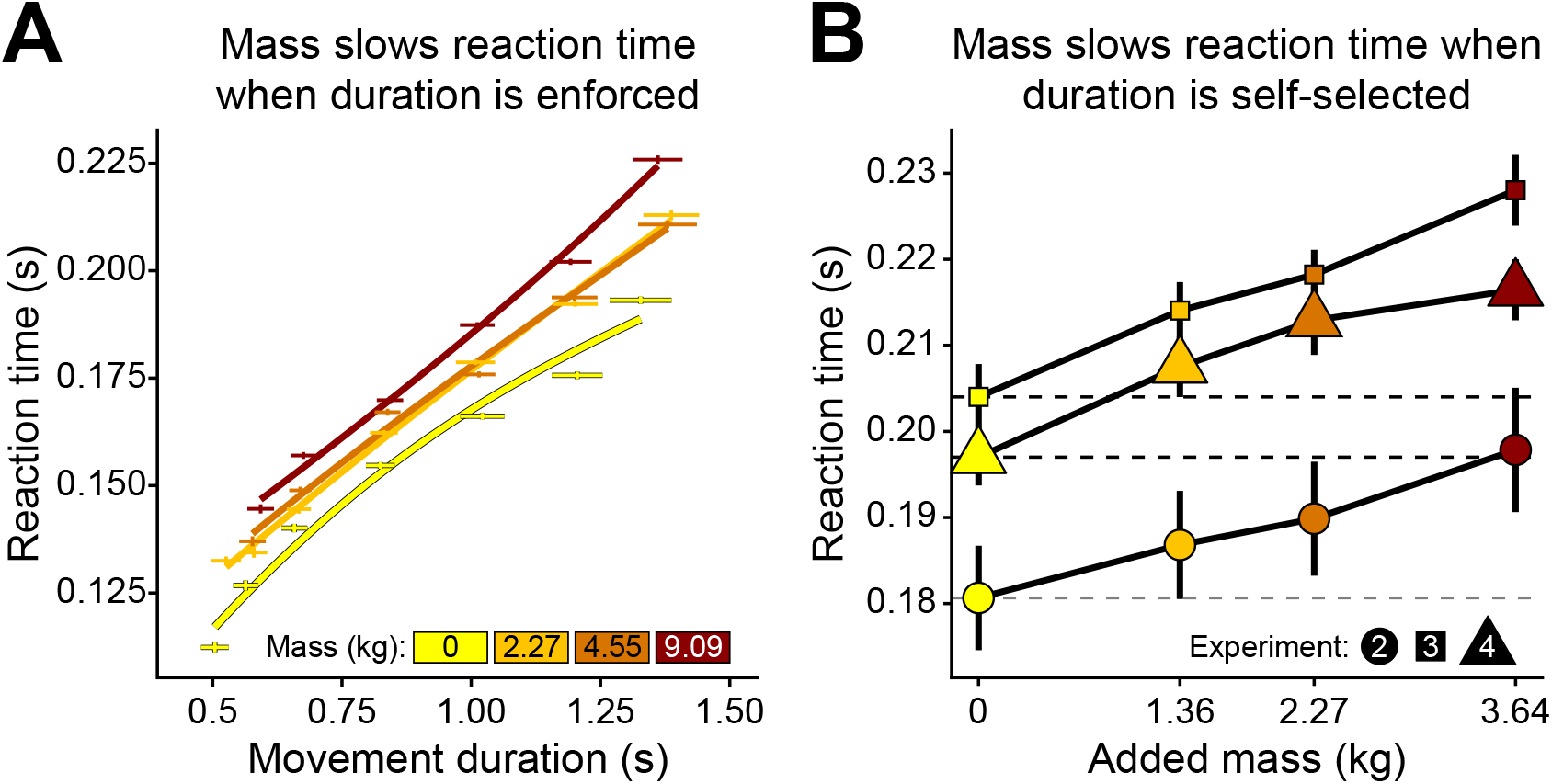
Reaction time findings. (A) Experiment 1 reaction times. At any given enforced movement duration, reaction times increased with added mass. (B) Experiments 2-4 reaction times. When movement durations were freely self-selected, reaction times increased with added mass across all accuracy conditions. The quickest reaction times were seen in experiment 2 (medium accuracy), then followed by experiment 4 (no accuracy), and lastly by experiment 3 (high accuracy).

## Discussion

The goal of this study was to understand how effort, represented as metabolic cost and modulated by mass, affects movement speed in arm reaching. We found that adding mass at the hand increased the metabolic cost of reaching and led subjects to make slower movements. However, when comparing the speeds that minimized metabolic costs to the empirical preferred reach duration of subjects, we found that individuals moved more slowly than what was predicted by minimizing metabolic cost alone. Incorporating an accuracy penalty to the proposed utility framework, which represents a reward-effort tradeoff discounted by time, more effectively explained preferred movement durations Shadmehr et al. [2016]. Altogether, these results suggest that effort is accounted for when determining the speed of reaching movements, but that other factors, namely accuracy, are incorporated in tandem.

### Utility predictions

The novelty of this study is the use of a single model to predict reaching duration across varying effort and accuracy constraints. Previous investigations have focused on these independently, but not in combination.

To complete a movement, a short planning phase exists during the reaction time period prior to movement initiation. To more accurately quantify the total cost of a movement, we need to account for this cost of reacting, or waiting. The added term *ė*_*r*_*t*_*r*_ accounts for the energy used while subjects are in the chair, planning the movement, but not yet moving. Adding this term aids the utility prediction by accounting for the effort expended during this planning phase, proportional to their resting metabolic rate. A future study may be able to determine an optimal reaction time and movement time that maximizes this utility.

The second addition to utility is a probability of reward based on movement vigor. This addition is similar to a speed accuracy trade-off, where faster movements tend to become less accurate Fitts [1992]. The opposite is also true: when accuracy requirements becomes more strict (harder to stop in target) movements slow down to maintain performance. Previous work has shown that depending on reach distance, variability of the movement changes O’Brien and Ahmed [2013]. However, this study did not investigate how movement variability affects self-selected speeds, only the extent of movement excursion. By including a probability of reward that is dependent on movement speed and mass into utility, we are able to reasonably predict the self-selected movement speeds across two experiments with a single *α* parameter. To our knowledge, this is the first study that has been able to reconcile multiple self-selected speeds within a single model.

### Effort and slowness

One common method used to explain movement speed is to minimize the total metabolic cost of the movement. Minimizing metabolic cost of transport has been used to explain preferred walking and running velocity in many paradigms Agiovlasitis et al. [2011], Bertram and Ruina [2001], Browning and Kram [2005], Handford and Srinivasan [2014], Hogberg [1952], Long and Srinivasan [2013], Minetti et al. [2003], Ralston [1958], Rathkey and Wall-Scheffler [2017], Seethapathi and Srinivasan [2015], Selinger et al. [2015]. While a convenient measure, cost of transport may not fully explain observed locomotion slowing in humans when carrying greater loads Hughes and Goldman [1970], as the metabolically optimal speed does not always shift with increasing mass Bastien et al. [2005]. In our study, we found that the metabolic minimum slows with increasing mass, but that preferred durations were slower than predicted by these metabolic minima.

An alternative explanation as to why we slow down with increasing effort is that for every movement there is only a finite amount of energy set aside for utilization, or an energetic budget, and thus we slow down with increasing effort to avoid exceeding this budget Hughes and Goldman [1970]. When walking with loads, Hughes and Goldman showed that the cost of walking (per kilogram meter) with added loads does not change. This may indicate that for a specific distance, humans have a specific energetic budget for the movement. When the effort of a movement increases and to keep the energetic expenditure within budget, we must slow down. Using the metabolic model (equation 3) and applying the masses and movement durations from experiment 2, we would find that the metabolic cost of the movements were 86.9, 89.5, 90.9, and 93.1 J for the 0, 1.36, 2.27, and 3.64kg conditions. If we normalize these values to the added mass and movement distance to arrive at comparable units as Goldman et al., we find 347, 226, 186, and 148 J kg^-1^ m^-1^. As effort increased, not only did movements slow, but the energetic budget changed.

Interestingly, we saw that the change in the metabolic minima is also very similar to the change in self-selected movement speeds. According to our metabolic model, the movement durations predicted by minimizing gross metabolic cost increased by 6.4%, 9.7%, and 13.8% with the masses seen in experiment 2 as compared to 0 kg (table 1). Empirical preferred durations from experiment 2 increased by 8.6%, 12%, and 16% for the added masses. All normalized movement durations are shown in table 1. Across experiments and metabolic optimizations, the normalized durations remain consistent. While metabolic cost may not alone predict movement vigor, it still was an important factor in determining changes in movement vigor.

### Accuracy

We predicted the discrepancies between metabolic minimum predictions and empirical preferred movement durations may have been due to an additional cost of accuracy. The speed-accuracy tradeoff predicts that smaller targets should incur a reduction in movement speed to sustain performance and, additionally, that slower movements are more accurate than fast movements when moving toward a target of the same size Dean et al. [2007], Fitts [1992]. We found that, indeed, slower movement durations increased reach accuracy across all experiments and when the accuracy requirements were altered, preferred reach durations adjusted accordingly (Fig. 3).

We also found evidence for an effect of mass on accuracy. In experiment 1 there was an effect of mass on endpoint error (i.e., accuracy) such that heavier loads worsened (shifted to the right) the speed-accuracy curve (Fig. **??**. However, during experiments 2 and 3 in which participants were free to self-select movement speed, we found no effects of mass on endpoint error. Thus, the effect of mass on reach accuracy found during experiment 1 may have been driven by enforcing durations and, across all experiments, endpoint error was more dependent on movement duration than mass. Mass also did not change the variability in endpoint error in any experiments, but as expected, slower movement durations reduced this variability. There was a small effect of mass on angular endpoint error variability (p = 0.0114), but this may be driven by the inertial properties of the arm. The lowest inertial directions of the arm are not oriented exactly at 45^*o*^ and 225^*o*^, which perhaps attracted reach trajectories to be slightly off center from the targets in order to adhere more closely to the axes of this inertial ellipse Goble et al. [2007]. Our findings critically implicate that both reach effort and accuracy are considered simultaneously when planning a goal-directed reach.

### Reaction Time

Our results show that imminent effort affects the reaction time of the movement. Across all our experiments, reaction time was self-selected by the subjects (including experiment 1) but consistently slowed with the prospect of added mass. This result mirrors earlier work on upper-limb movements showing that an increase in required distance Rosenbaum [1980], Reppert et al. [2018] or isometric force production Stelmach and Worringham [1988], Ivry [1986] leads to a lengthening of reaction time or latency of the action. While the slowing of reaction time has generally been explained as a consequence of increased evidence integration, usually due to increased uncertainty Mormann et al. [2010], Ratcliff and Van Dongen [2011], it is possible that a general decision variable that tracks utility of a movement is being integrated over this duration. Therefore, a lower utility movement that takes longer to integrate a decision variable to a threshold could explain the slowing of reaction time Shadmehr and Ahmed 2020].

An alternative theory postulates that the speed and latency of actions is inherently affected by the opportunity cost of time Niv et al. [2007], Yoon et al. [2018], which is proportional to the rate of reward/utility of the environment in which the movements are being made. In other words, the utility of a movement is not just determined by immediate factors but also by the past experience and future expectations of an environment. A more effortful environment therefore has a lower opportunity cost of time, owing to lower net utility rate, and thereby promoting more slothful reaction and movement times. Further evidence of this is seen in experiment 1 wherein subjects react slowly not just to added mass but also to increases in enforced movement duration (Fig. 4A).

The duration between stimulus onset and initiation of a movement encompasses cognitive, neural and physiological processes that result in a movement being successfully initiated and completed. While theories of response or reaction time have focused on cognitive factors and corresponding neural implementation, the (neuro–)physiological aspect of preparing a movement has only been recently considered. An increase in required effort could increase the duration required to prepare or “organize” the movements Nagasaki et al. [1983] and generate the appropriate force Irie et al. [1983]. Further research is required to disambiguate the various possible effects of added effort on reaction time through a series of carefully controlled behavioral experiments.

### Limitations and Future Work

While our model of movement vigor was able to adequately adjust across contexts, there were still many limitations introduced during its construction. First, we estimated speed-accuracy tradeoff curves from experiment 1, which was a separate task with somewhat differing goals. While accuracy was required and encouraged in experiment 1 in order to receive feedback regarding the enforced reach duration, accuracy itself did not carry any penalty to task performance. In other words, missing the target in experiment 1 did not result in a loss of reward, points, or failure. Thus, it is possible that generalizing experiment 1 endpoint errors to estimate accuracy in experiments 2 and 3 may have introduced error. Along this line, we opted to exclude modeling of experiment 4 due to its out-and-back nature differing from the out-and-stop paradigm of the other experiments.

Second, the “reward” in our experiments was of nebulous value and was fit as a free parameter in our model. We opted to increase model generalizability by fitting reward value (*α*) from experiment 2 data, and then using the same *α* in predicting experiment 3. Regardless, it is possible that some of the incongruities between the metabolically predicted preferred durations and the empirical durations could have been due to inadequately appraising the reward, as humans and nonhuman primates accelerate movement speeds when expecting rewards Milstein and Dorris [2007], Summerside et al. [2018], Takikawa et al. [2002], Xu-Wilson et al. [2009]. Individuals opted to move more slowly than predicted by minimizing effort alone and we assumed this was due to the cost of accuracy; however, it is also possible that the value assigned to these movements was lower than predicted.

Third, the model presented here does not differentiate between the objective, metabolic effort of movement and the subjective perception of this effort. During our paradigms, we used a constant set of masses across all individuals, rather than scaling the applied mass based on body mass or strength. Participants may have perceived the effort of moving, for example, the 2.27 kg mass differently dependent upon strength, dopamine levels, or even fatigue Reppert et al. [2018], Shadmehr et al. [2016], Goh et al. [2021], Chong et al. [2017]. Again, observed differences in predicted and actual movement durations could also be the result of individual differences in effort sensitivity.

And lastly, our reward rate model accounts for both reaction and movement times. We used measured reaction times from experiments 2 and 3, and only solved for movement time. Future iterations of this model should seek to optimize for both reaction and movement times. Successfully modeling reaction times may provide valuable information regarding how utility of a movement impacts this planning phase and may offer an additional parameter that an individual can tune when adjusting their strategy Haith et al. [2016].

## Conclusion

In this study we examined the effect of effort on the metabolic cost and the movement duration of reaching movements, and then modeled this movement speed using metabolic cost or a utility framework. Added effort (mass), lead to an increase in metabolic rate and cost. An effort model (equation 1) predicted that this increase in cost is sublinear with mass. Adding mass (effort) to a movement also increased the preferred movement duration, but exclusively minimizing the metabolic cost of a movement was not able to predict preferred durations. Instead, using a utility framework that incorporated accuracy alongside effort (equation 5), we were able to reasonably predict the observed movement durations within distinct environments. Collectively, our results provide novel evidence for the joint importance of reward, accuracy, and effort, as represented by metabolic cost, on reaching vigor. Our work here lays fundamental knowledge regarding the metabolic costs of arm reaching movements, and sets the stage for systematically changing effort based on metabolic cost in future paradigms.

## Methods

This study is composed of four experiments and a model analysis. The first experiment measured the effect of mass and speed on the metabolic power of reaching. The remaining three experiments determined how mass and accuracy coalesce to influence preferred reaching speed. In the model analysis, we used the measured metabolic data from experiment 1 to determine which utility formulation can best explain the preferred movements observed in experiments 2 and 3.

### Experiment 1 - Effect of mass on metabolic power

In the first experiment, 5 male and 3 female, all right-handed, with an average age of 28.9 years (std = 5.5), average weight 66.7 kg (std = 11.7), and average height 173.4 cm (std = 10.4) completed the protocol. All subjects except one completed the experiment in two sessions. The remaining subject completed the protocol over 3 sessions. All participants reported no neurological, cardiovascular, or biomechanical problems that could interfere with the study. Subjects gave written informed consent, as approved by the University of Colorado Institutional Review Board.

#### Protocol

Subjects completed reaching movements with varying speed and mass requirements. Reach kinematic and metabolic data was collected as a function of mass and speed (Fig. 1A). Subjects sat in a chair that was height adjusted to place the screen 3 feet in front and 1 foot above of the subjects’ line of vision, with their arm in a horizontal planar position. They were trained to move a cursor from a home circle and stop at a target circle within a specified time window. Subjects made reaching movements in seven distinct time windows across four different masses (Fig. 1A). A block refers to one speed combined with one mass condition. The number of trials per block was determined such that each block consisted of five minutes of reaching at the desired speed, where the first 20 trials of each block were used for training. To begin a trial subjects held a circular cursor (r = 0.4 cm, yellow colored) within the home circle (r = 1.1 cm, white circle) location for 200 ms. The home circle then disappeared and a target circle 10 cm away (r = 1.4 cm) appeared randomly at 45, 135, 225, and 315 degrees from the right horizontal. In training, a blue dot would make a simulated movement from the home circle to the target circle using a minimum jerk trajectory. Feedback on movement duration was given when the center of the cursor was within the target the first time. If subjects moved too slow the target circle would turn grey, whereas if the subject moved too quickly the target would turn green. Appropriately timed movements resulted in the target flashing yellow and a pleasant tone. Upon completing an outward reaching trial, the home and target circle would swap locations and the subject would make another reaching movement towards the center of the screen. Subjects completed four different mass conditions and six different speeds. The completed mass conditions were 0 kg (0 lbs), 2.3 kg (5 lbs), 4.5 kg (10 lbs), and 9.1 kg (20 lbs) of added mass at the robot handle which supported the vertical mass. The seven different time windows were: Very, Very Slow (VVS, 1.25 – 1.35 s, 160 trials), Very Slow (VS, 1.05 – 1.15, 170 trials), Slow (S, 0.85-0.95 s, 200 trials), Medium (M, 0.65-0.75 s, 220 trials), Fast (F, 0.45-0.55 s, 240 trials), Very Fast (VF, 0.325-0.425 s, 250 trials), and Very, Very Fast (VVF, 0.225-0.275 s, 260 trials). For 0 kg and 2.3 kg added, subjects would complete the faster speed conditions of VS to VVF. For 4.5 kg and 9.1 kg added, subjects would complete the slower speed conditions of VVS to VF.

#### Metabolic Data Collection

Metabolic data was collected for the duration of each five minute reaching block. Subjects wore a nose clip and breathed into a mouthpiece, connected to a metabolic cart (ParvoMedics, TrueOne 2400), which measured 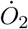 consumption and *C*?*O*_2_” production. Subjects were required to be well rested and have fasted for 8 hours before testing. Testing sessions began with the subject resting in a seated position in a chair for 10 minutes. Three baseline readings were then taken for 5 minutes each before the experimental protocol began. During these baseline readings, subjects sat quietly in the experiment chair, and held the robotic arm manipulandum. Subjects then began the arm reaching trials. Five-minute rest periods were provided between each block of reaching trials.

### Experiments 2, 3, & 4 - Effect of mass on preferred movement duration

In the second and third experiments, a separate cohort made seated horizontal arm reaching movements using a robotic arm manipulandum (Interactive Motion Technologies Shoulder-Elbow Robot 2) while secured to a chair by a 4-point seat belt. The seat was height-adjusted to place the screen 3 feet in front and 1 foot above the participant’s plane of view, with their arm in a horizontal planar position in a similar manner to experiment 1 (Fig 1A). All participants provided written informed consent and reported no neurological, cardiovascular, or biomechanical problems that could interfere with the study.

#### Kinematic Data Collection

Subjects made reaching movements within five blocks of mass conditions. The five blocks were a familiarization block, 0 kg, 1.36 kg (3 lbs), 2.27 kg (5 lbs) or 3.64 kg (8 lbs) added at the hand. The order of the weighted conditions was randomized for each subject. The downward weight of the added masses was supported by the robot, so these masses only added inertial effects to the arm. The position of the handle controlled a cursor on a computer screen that was placed just above head level and about 3 feet in front of the subject. Subjects arm positions started in approximately the same orientation. Visual feedback was provided to the subjects throughout the experiment on whether they completed the reach movement in the prescribed duration. To begin a trial subjects held a circular cursor (r = 0.4 cm, yellow colored) within the home circle (r = 1.1 cm, white circle) location for 200 ms. The home circle then disappeared and a target circle (shape dependent on experiment) 10 cm away appeared randomly at 45, 135, 225, and 315 degrees from the right horizontal. A subject would go through all four outward targets in a pseudorandom order then begin again (Fig. 1B).

*Experiment 2*. In this experiment 8 male and 4 female subjects, all right-handed, with an average age of 26.2 years (std = 3.1), average weight 68.4 kg (std = 4.4), and an average height of 173.6 cm (std = 11.1) completed the experiment (Fig. 1B). Subjects underwent 5 different blocks of 400 reaching movements (200 out and back movements) to four different targets. The subjects would make horizontal arm reaching movements towards a circular target, similar to experiment 1 (r = 1.4 cm, red color). For each reaching movement, the target would explode indicating a correct movement duration as the movement duration criteria was set between 1ms and 10000 ms, so there were not time requirements imposed on the reaching movements. In this experiment we wanted to ensure subjects came to a complete stop before the trial ended, so the dot would explode after the subject had remained in the target for 300 ms and the velocity during that time was under 0.5 mm/s.

*Experiment 3*. We wanted to ensure that the effects on movement duration from mass were not influenced by the size of the target or accuracy costs. We ran two more similar experiments in which we altered the shape of the target to change the accuracy costs of the movement.

In experiment 3, there were 9 male and 3 female subjects who completed the protocol, with an average age of 25.0 years (std = 3.61), average height of 171.5 cm (std = 7.58), and an average weight of 67.5 kg (std = 2.99). Subjects completed 400 out and stop trials in familiarization, and 200 out and stop for each added mass condition. In this experiment, the target was a section of a 10 cm circular arc centered on the home circle oriented at the same positions as experiment 1 and 2. Subjects would need to stop between 10 cm and 11 cm from the home circle and within 7 degrees (2.44 cm arc length) of the center of the arc target for the target arc to turn green. If subjects overshot the target (went past 11 cm) the target would turn red indicating an overshoot. Thus, this experiment had tighter accuracy constraints in the radial direction (1cm) compared to experiment 2 where the equivalent constraint in the radial direction was 2.8 cm.

*Experiment 4*. Experiment 4 was completed by 18 subjects, 9 male and 9 female, with an average age of 25.1 ± 3.7 years old. Subjects completed 100 out and back movements for familiarization, and 200 out and back movements for each mass condition. The target was a 90-degree section of a circle that subjects had to reach towards. However, subjects did not need to stop at any specific location, just hit the target arc, turn around and return to the home circle. In this experiment we used the point that they turned around as the end of their movement. This experiment was used to simulate a zero-accuracy cost with a very large target.

### Data Acquisition and Analysis

For all experiments, robot handle X and Y position data was recorded at 200 Hz and analyzed in MATLAB 2019a. Position data was filtered using a fourth order low-pass Butterworth filter (cutoff frequency 10 Hz) and differentiated using a double five-point differentiation to obtain velocity and acceleration. To calculate radial velocity, we calculated the Euclidean distance from the home circle and differentiated using five-point differentiation.

#### Metabolic Processing

The gross metabolic rate was calculated in joules per second, *ė*, using the method described by Brockway (Eq. 7) (Brockway, 1987):

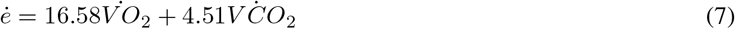

Average baseline metabolic rate from the three baseline sessions was subtracted from gross metabolic rate to determine the metabolic rate associated with the reaching movement only, or net metabolic rate. Subjects made reaching movements for 5 total minutes, but metabolic rate was calculated only using the last 3 minutes of each block to allow subjects to reach a steady metabolic rate while reaching. After data was collected, custom MATLAB scripts were used to parse the data by trial, mass, and speed. Movement duration was calculated using the last 3 minutes within a block. The overall metabolic rate is then normalized by the fraction of time spent moving.

#### Metabolic cost models

We used the measured metabolic power data, *ė*, to parameterize the metabolic cost of a movement, *E*, as a function of mass and movement duration. Parameter estimates were computed using the function nls from package nlstools. Using the data in experiment 1, we fit gross metabolic power to the following functiom (equation 8) using average subject effective mass (described later in methods), metabolic power and movement durations. The fitted parameters are *a, b, i*, and *j*, where *a* represents is an offset representing the cost of not moving, *b* is a scaling parameter, *i* shows how effort scales with mass, and *j* shows how effort scales with time. Further, m represents the effective mass of the movement and *T* represents the movement duration.

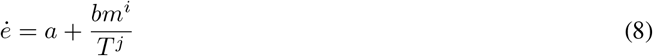

To obtain an expression for the total metabolic cost of the movement, *E*, we multiply equation 8 by movement duration, T:

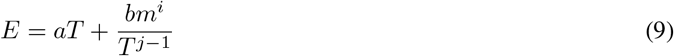

Importantly, the above expression also tells us the metabolically optimal movement duration for a movement of a given mass.

To determine whether the cost of resting was influenced by added mass, an additional metabolic cost model was fit to gross metabolic power data that included a term for effective mass that multiplies the *a* parameter. This represented a mass-dependent cost of resting, where *k* is the scaling on the resting mass cost:

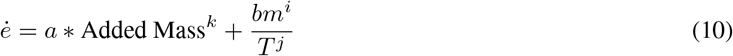

AIC and BIC scores are calculated for each metabolic cost model.

#### Movement Onset and Offset

Many of our metrics are dependent on identifying the instant the movement began (movement onset) and the instant the movement ended (movement offset). To detect movement onset and offset we used a custom algorithm for all experiments. Movement onset algorithms are often influenced by movement speed and we wanted to minimize this effect Brenner and Smeets [2018], Reppert et al. [2018]. We first computed the velocity towards the target by differentiating the distance from the center of the target. For movement onset we find the first time the velocity towards the target reached 20% of the maximum velocity towards the target. From there the algorithm searches backwards (in time) to a point where either there is either 4 frames of acceleration away from the target in the next 10 frames or where the standard deviation in the velocity towards the target was less than 2e-3 in those 10 frames. Using this method, we detect the first time the subject begins consistently accelerating towards the target. This led to detecting movement onset much earlier than many velocity thresholding algorithms, where we find an average velocity at movement onset of -0.389 ± 8.019for experiment 1 (mean ± sd), 1.019 ± 2.220 mm/s in experiment 2, -0.052 ± 5.552 m/s for experiment 3, and -0.098 ± 5.077 mm/s in experiment 4. To detect movement offset in experiment 2 and 3, we found the first time the reaching movement was first 9 cm away from the home circle, then used the same method as reaction time to determine the offset. We find the first time the standard deviation of the velocity towards the target is less than 2e-3. In experiment 4, movement offset is determined as the point subjects turn around and begin moving back towards the home circle.

Because reaction time algorithms can be dependent on movement duration Brenner and Smeets [2018], Reppert et al. [2018], we needed to compare these changes in reaction time to computed reaction times from simulated movements. These movements can be generated using the range of movement durations in experiment 1 or 2. This will inform us if the reaction time changes are due to changes in movement duration or added mass. We simulate reaching movements with similar movement times to the experiment using a minimum jerk trajectory and a simulation of the arm making reaching movements Flash and Hogan [1985]. Using the minimum jerk simulation, we found that over the span of movement durations for experiment 1 the reaction time would increase with increasing movement duration. Over the movement duration ranges (.45 s – 1.4 s) the reaction time would increase due to the algorithm by 40 ms. Simulating the minimum jerk for experiment two, over the movement durations (0.77s – 0.90s), the reaction time would increase by 0.3 ms. We also used a biomechanical model of the arm like other studies to test if the change in reaction time could be attributed to subjects using the same control signal for different masses Li and Todorov [2007]. The reaction time change given the same control signal across masses was small, about 3 ms. In experiment one, the average reaction times across speeds ranged from 0.1377s to 0.245s. This is much larger than the effect of movement duration from the simulated movement durations simulation like experiment one (108 ms experimental vs 40 ms simulated). The algorithm used is affected by movement duration, but the change in calculated reaction time is greater than the range from just the effect of speed on the calculated duration.

In experiments 2 and 3, the range of reaction times calculated from the simulated movements was 0.3 ms. The reaction times calculated from this experiment ranged by 17 ms for experiment 2, 24ms in experiment 3, and 15 ms in 4. The reaction time range in experiment one has a much larger range than the calculated reaction times of the simulated movements. This indicates that the changes in reaction time were due to experimental manipulations of mass, not the movement onset algorithm or changes in the movement durations.

#### Movement Duration

Movement duration is calculated as the time between movement onset and movement offset, except for experiment 4 where it is the time between movement onset and when the cursor crosses the 10 cm perimeter surrounding the home circle. The realized movement times in experiment 1 were generally longer than the prescribed movement times in the protocol. This is due to the feedback being given before the end the movement as subjects would take some time to settle on the target and feedback was given as soon as they reached the target.

#### Error Calculations

We investigated three measures of error for all experiments. Endpoint error was the Euclidean distance between the cursor at movement offset and the center of the target. We then broke endpoint error into two components, angular error, and radial error. Angular error was calculated as the angle between the vector pointing from home circle to target circle, and the vector from home circle to the cursor at movement offset. A clockwise angular error was considered negative. The second metric, radial error, was calculated as the distance from cursor to the target center (10 cm from the home circle, center of the arc) at movement offset along the radial axis. Maximum excursion was also calculated as the greatest Euclidean distance from the home circle during the movement. Experiment 4 had no stopping criteria, so endpoint error is defined as the maximum excursion.

#### Outlier Analysis

In experiment one, we removed outlier trials from the statistical analysis if they did not complete the movement correctly. Movement kinematic metrics were computed for every trial, then we removed trials based on specific criteria. We removed any trial where the endpoint error was greater than 10 cm (reached the wrong target), the movement duration was less than 0.2 seconds or greater than 2 seconds (did not make the movement), the reaction time was greater than 0.50 s (failed to initiate movement), or the absolute angular error is greater than 50 degrees (reached to wrong target).

For the kinematic statistical analysis, the data is then split into outward and inward reaching movements and we only use the outward reaches in the statistics. Inward and outward trials are split because on inward movements subjects knew where the target would show before it was indicated, which affects movement kinematics. After removal of trials, 27 out of 15925 outward trials were removed.

In experiments 2, 3, and 4, we removed trials that were outside 1.5x the interquartile range of movement duration, reaction time, reaction velocity, or angular error Ghasemi and Zahediasl [2012]. Reaches with a maximum excursion of more than 14cm were also filtered out in experiment 2 and 3. The data from experiment 2 and 3 were also split into outward and inward trials for the same reason as experiment 1. Statistical and kinematic analysis were done on the outward trials. This removed 702 out of 9600 outward trials in 2; 671 trials of 9600 are filtered out for experiment 3; 2335 of 14400 trials are filtered out for experiment 4.

### Effective Mass Calculation

For all models, mass was represented by the subject-specific effective mass of the arm, averaged over the four reach directions. Effective mass is an estimate of the arm’s resistance to motion when to a force applied in each direction Shadmehr et al. [2016]. Segment lengths and masses are estimated from anthropometric tables. Contini [1972], Enoka [2002], Winter [2009].

To determine the effective mass of the arm at a given time point we defined the Jacobian matrix for a two-link model of the arm, Λ, where *l*_1_ is the length of the upper arm and *l*_2_ is the length of the forearm. *θ*_*s*_ and *θ*_*e*_ are the shoulder and elbow joint angle respectively:

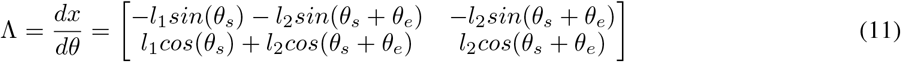

The inertial matrix (*I*(*θ*)) is defined in Eq. 12, where mass is mass added at the hand. The centroid lengths, *r*_1_ and *r*_22_, refer to the centroid length of the upper arm and forearm with mass added. *I*_*COM*,1_ and *I*_*COM*,2_ are the moment of inertia about the center of mass for the upper arm and forearm.

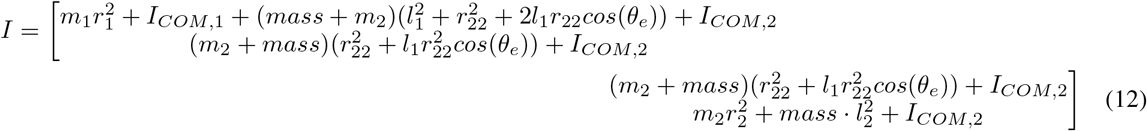

The mass matrix (*M*) is defined as:

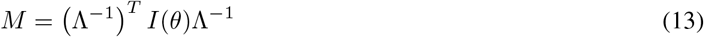

We obtain the effective mass, m, in a given reach direction by applying a unit vector acceleration in that direction and calculating the magnitude of the resultant force vector. Each subject’s specific effective mass for experiment 1 and 2 was calculated using anthropomorphic measurements and estimates Contini [1972], Huston [2008], Winter [2009]. In the first experiment the average effective masses of the four mass conditions were 2.44±0.064, 4.834±0.068, 7.127±0.070, and 11.691±0.071 kg. In experiment 2 the effective masses of the subjects and robot arm in the four mass conditions were 2.506±0.073, 3.959±0.073, 4.894±0.075, and 6.282±0.076 kg. The average effective mass in experiment 2 is used for effective mass in experiment 3.

### Models of optimal movement duration

Here we describe the modeling analysis employed to calculate predicted optimal movement durations with changing effort and accuracy requirements. We use the group average data for effective mass, accuracy, movement duration, and reaction time.

#### Optimal duration based on maximizing net reward rate

When the goal of the movement is maximizing net reward rate, the utility is determined by the sum of the reward of the movement (*α*) minus the sum of the effort (*E*), both discounted by time (*T*) and is shown in Eq. 14.

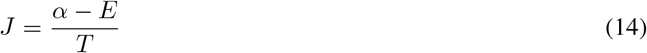

Effort is the metabolic cost of the movement (Eq. 15) and is represented by the expression obtained from the measured metabolic data in experiment 1 (Eq. 8). Total time is split into reaction time, *t*_*r*_, and movement time, *t*_*m*_. We use the measured reaction times from experiments 2 and 3 and solve for the movement durations that maximize utility. Here the only parameter to be fit is *α*, which represents the subjective reward associated with completing the arm reaching movement. The reaction times used from experiment 2 are 0.178 ± 0.013, 0.185 ± 0.014, 0.190 ± 0.013, 0.196 ± 0.013 for 2.47kg, 3.80 kg, 4.70 kg, 6.10 kg respectively. Experiment 3 reaction times were 0.205 ± 0.013, 0.215 ± 0.013, 0.219 ± 0.013, 0.229 ± 0.013. Experiment 4 reaction times were 0.199 ± 0.012, 0.208 ± 0.011, 0.213 ± 0.012, 0.216 ± 0.012.

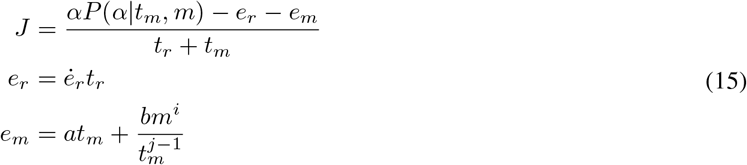

A single parameter alpha is used to fit data from experiment 2 and used to predict the durations in experiment 3. The main difference between experiment 2 and 3 is the size of the target, which can be represented as the probability of acquiring reward at a given movement duration. To incorporate this speed-accuracy tradeoff into the utility equation, we scale *α* by the probability of stopping within the target given the movement duration and mass:

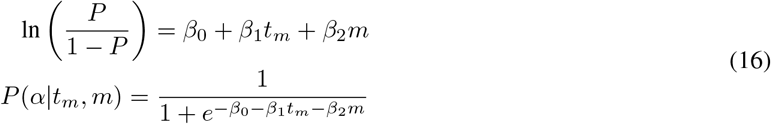

The probability function scaling *α* was determined using target criteria from experiment 2 and 3 and movement kinematic data from experiment 1. Each arm reaching movement in experiment 1 was labeled as a success according to the success criteria (target size) from experiment 2 and 3. Thus, each trial in experiment 1 will have two labels for success. The first label from success in experiment 2 and the second from success in experiment 3. For experiment 2 labeling, we labeled a reach as a success if the endpoint error was less than 1.4 cm. For experiment 3, we labeled each reach as a success if the maximum excursion was less than 11 cm (and greater than 10cm), and the angular error was less than 7 degrees. After determining if each reach was a success or not, we use an inverse logistic regression (R, glm model with binomial family and logit link function) to fit function *P* (*α*|*t*_*m*_, *m*) for each experiment, obtain two sets of beta coefficients.

For experiment 2, we found the beta coefficients of the regression to be *β*_1_ = -0.0946±0.008 for mass, *β*_2_ = 5.877±0.188 for movement duration, and an intercept of *β*_0_ = -1.450±0.125. Experiment 3 had coefficients of *β*_1_ = -0.0828±0.006 for mass, *β*_2_ = 6.0912±0.129 for movement duration, and *β*_0_ = -2.9312±0.091 for an intercept. These logistic regressions represent *P* (*α*|*t*_*m*_, *m*) in the utility model.

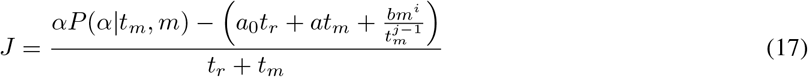

We optimize this utility model (Eq. 17) by finding the value that minimizes the sum of squared error between predicted movement duration and the average movement duration from experiments 2. Data was analyzed with the function optimize in R and a custom written error function.

#### Optimal duration based on minimizing metabolic cost

The metabolically optimal duration is the movement duration that minimizes the metabolic cost of movement of a given mass. This is calculated, as described earlier, by minimizing Eq. 9.

### Statistical Tests

Kinematic data was exported to R (v 1.2.5001) for statistical analysis. Linear mixed effects models were computed using the lme4 and multcomp package, and the functions used were lmer and cftest. To analyze the effect of mass on the kinematic variables we used linear mixed effects models. The effective mass used in the linear mixed effects models is the average effective mass from experiment 1 and experiment 2. Experiment 2 effective mass values are applied to experiment 3.

For experiment 1, we tested a model with no between-subject variables and added mass and movement duration as within-subject variables. Gross metabolic power and reaction time are the dependent variables.

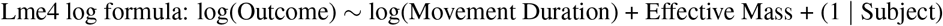

For experiment 2, 3 and 4 we tested the main effect of added mass and movement direction on movement duration, peak velocity, and reaction time.

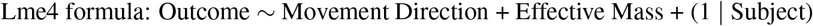

An ANOVA (aov in R) with post-hoc Tukey tests (TukeyHSD) was used to test differences between experimental outcomes (2, 3, and 4). Experimental data was aggregated for each experiment and subject before use in the ANOVA and Tukey test.

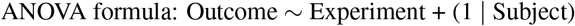

We use a significance level of *α* = 0.001351 or 1.351e-3 as we made 37 comparisons. We compute 29 linear mixed effects models and 2 ANOVAs with 3 Tukey Post Hoc comparisons for each ANOVA. Exact p-values are reported unless it is less than 2e-16. The linear model estimates are reported for the significant variables. For non-significant factors only the p-value is reported.

